# Extensive individual differences of category information in ventral temporal cortex in the congenitally blind

**DOI:** 10.1101/2020.06.14.151092

**Authors:** M. Rosenke, J. Van den Hurk, E. Margalit, H. P. Op de Beeck, K. Grill-Spector, K. S. Weiner

## Abstract

Human ventral temporal cortex (VTC) is a cortical expanse that performs different functions and computations, but is especially critical for visual categorization. Nevertheless, accumulating evidence shows that category-selective regions persist in VTC in the absence of visual experience – for example, in congenitally blind (CB) participants. Despite this evidence, a large body of previous work comparing functional representations in VTC between sighted and CB participants performed univariate analyses at the group level, which assume a homogeneous population – an assumption that has not been formally tested until the present study. Specifically, using fMRI in CB and sighted participants (male and female), we empirically show that at the group level, distributed category representations in VTC are more reliable in the sighted (when viewing visual stimuli) compared to the CB (when hearing auditorily-substituted visual stimuli). Despite these group differences, there is extensive heterogeneity in VTC category representations in the CB to the point that VTC category representations in a subset of CB participants (some who were born without eyes, but not all) are more similar to sighted individuals compared to other CB participants. Together, our findings support a novel idea that driving factors contributing to the formation of VTC category representations in the blind are subject-specific, which complements factors that may generalize across group members. More broadly, the present findings caution conclusions of homogeneity across subjects within a group when performing group neuroimaging analyses without explicitly quantifying individual differences.

## Introduction

Ventral temporal cortex (VTC) is a large cortical expanse involved in different functions, but is especially crucial for the neural processing underlying visual categorization (Grill-Spector and Weiner, 2014). For example, VTC contains both clustered (Kanwisher et al., 1997; Epstein and Kanwisher, 1998; Cohen et al., 2000; Downing et al., 2001) and distributed (Haxby, 2001; Spiridon and Kanwisher, 2002; Cox and Savoy, 2003; Kriegeskorte et al., 2008; Sayres and Grill-Spector, 2008; Walther et al., 2009; Weiner and Grill-Spector, 2010; Kravitz et al., 2011; Jacques et al., 2016; Grill-Spector et al., 2017) representations of object categories. Accumulating evidence from previous studies has shown that category-selective regions persist in VTC in the absence of visual experience. For example, a large-scale map distinguishing animate from inanimate representations in lateral and medial VTC is present in blind participants (Mahon et al., 2009), as are smaller clustered regions selective for objects (Amedi et al., 2007; Petrini et al., 2014), scripts (Reich et al., 2011; Striem-Amit et al., 2012a), tools (Peelen et al., 2014), body parts (Kitada et al., 2014; Striem-Amit and Amedi, 2014), non-manipulable large objects (He et al., 2013) and scenes (Wolbers et al., 2011).

Despite this evidence, nearly all previous studies performed univariate analyses at the group level (but see Peelen et al., 2014; van den Hurk et al., 2017). Group analyses assume a homogeneous population, which has never been formally tested. Recently, van den Hurk and colleagues showed that categorical information in VTC could be classified above chance in both sighted (audio and video stimuli) and blind (audio stimuli) subjects in which classification performance was higher for sighted compared to blind subjects. In the current paper, we build on this previous work and consider a *group hypothesis* and an *individual differences hypothesis* of distributed category selectivity in VTC of congenitally blind and sighted participants. A *group hypothesis* predicts that category representations will largely be homogenous among group members, but that the structure of these representations will differ between sighted and blind groups. An *individual differences hypothesis* predicts that irrespective of group-level differences, category representations are significantly more heterogenous among group members to the point that there are similarities between individuals across groups that are larger than similarities between individuals within a group. Empirically supporting either hypothesis has implications for future studies comparing populations at the group and individual level (Temple et al., 2001; Avidan et al., 2005; Duchaine and Nakayama, 2005; Cantlon et al., 2006; Golarai et al., 2007; Scherf et al., 2007a; Rykhlevskaia et al., 2009; Deen et al., 2016; Green et al., 2019b, 2019a).

We leveraged previously published data (van den Hurk et al., 2017), which contained functional and anatomical images from 14 congenitally blind and 20 sighted controls, to ask three main questions: (1) Are category representations in VTC similar between sighted and blind subjects at the group level? (2) Are individual differences in representational structures in VTC smaller within a group compared to between groups for both the blind and the sighted (in support of the *group hypothesis*)? (3) Despite any group differences, are there cases in which RSMs are more similar between subjects across groups (sighted and blind), compared to within group (in support of the *individual differences hypothesis*)?

## Materials and Methods

The Materials and Methods are similar to those published previously using this dataset (van den Hurk et al., 2017) up to the section *General Linear Model (GLM)*. We include the similar and new methods here to assure that the paper is self-contained.

### Subjects

Fourteen congenitally blind (5 females, mean age 37.1 years old) and twenty sighted (11 females, mean age 34.5 years old) adults with normal or corrected-to-normal vision took part in this study. We could not record auditory runs for two sighted subjects due to technical problems, resulting in 18 subjects with both visual and auditory data and 2 subjects with only visual data. Subjects were screened for fMRI compatibility, signed informed consent, and were financially compensated for their participation. The study was approved by the medical ethical committee of KU Leuven. Data from this study have also been reported in van den Hurk et al. (2017).

### Gradation of blindness

The blind subjects had to meet two inclusion criteria that are typical in the congenital blindness literature: 1) the onset of their blindness had to be congenital (from birth), and 2) they had no (memory of) object or shape perception. We assigned the blind volunteers to three categories of blindness. Category 1: anophthalmia (i.e. no eyeballs developed in utero). Category 2: no (memory of) light/dark perception. Category 3: light/dark perception in one or both eyes. See Table 1 in van den Hurk et al. (2017) for more details. Objectively, Category 1 blind subjects are the only category for which one can be certain that no visual perception was possible even in early post-natal weeks/months.

### Stimuli

We used four categories for both visual and auditory stimulation: faces, body parts, artificial objects, and scenes.

#### Visual

Visual stimuli consisted of 64 short (~1800 msec) movie clips with 16 unique clips per category. In the face category, clips portrayed actions such as laughing, chewing, blowing a kiss, and whistling. In the body parts category, clips showed scratching, hand clapping, finger snapping, bare feet walking, and knuckle cracking. In the object category, we created clips of a stationary car, washing machine tumbling, ticking clock, tools, mug, drinks poured in glass, bouncing ball, and rotating fan. In the scenes category, the clips depicted waves crashing on a beach, a busy restaurant overview without clear faces visible, a train station with departing trains, moving traffic, forest with rustling leaves, a lake with birds, and a grass field with moving grass.

#### Auditory

Audio stimuli consisted of 64 short audio clips (~1800 msec) with 16 unique clips per category, and were repeated across runs. Stimuli were matched in overall sound intensity by normalizing the root-mean-square of the sound pressure levels. Each sound stimulus was matched (as closely as possible) to each visual stimulus (i.e., the visual movie clip of clapping hands and the sound of clapping hands, etc).

### Experimental Design

For sighted subjects, we scanned 4 functional runs of each modality (auditory and visual), with the auditory runs presented first. For blind subjects, we scanned 8 functional auditory runs of which we used the first 4 runs for the present analyses in order to match the number of runs across groups and modalities. For both modalities and within each run, the stimuli were presented in a block design, with 4 blocks per condition per run. In each block, 8 stimuli (each ~1,800 msec duration) were presented with a 200 msec inter-stimulus interval. The presentation order of conditions was counterbalanced within and between runs to account for possible order effects. Each run lasted for approximately 7.5 minutes. For the remainder of the manuscript, *sighted visual* refers to fMRI data acquired in sighted subjects while viewing visual stimuli for 4 experimental runs, while *sighted audio* refers to fMRI data acquired in sighted subjects while listening to auditory stimuli for 4 experimental runs.

### MRI data acquisition parameters

Functional and anatomical images were acquired on a 3T Philips Ingenia CX scanner (Department of Radiology, University of Leuven) with a 32-channel head coil. Each functional run consisted of 225 T2*-weighted echoplanar images (EPIs) (1.875×1.875 mm in-plane voxel size, 32 2.2 mm slices, interslice gap 0 mm, TR = 2000 msec, TE = 30 msec, 112×112 matrix). In addition to the functional images, we collected a high-resolution T1-weighted anatomical scan for each subject (182 slices, resolution 0.98 by 0.98 by 1.2 mm, TR=9.6 msec, TE = 4.6 msec, 256×256 acquisition matrix). Stimuli were presented using custom written MATLAB R2014a code (Mathworks Inc, Natick, MA) and Psychtoolbox 3 (Brainard, 1997) via an NEC projector with an NP21LP lamp that projected the image on a screen the subject viewed through a mirror. Viewing distance was approximately 64 cm.

### Data preprocessing

Functional and anatomical data were pre-processed using the BrainVoyager QX2.8 package (Brain Innovation, Maastricht, The Netherlands), while the remainder of the data processing and analyses were performed using custom written MATLAB code in combination with the NeuroElf v0.9c toolbox (NeuroElf.net). Functional volumes were first corrected for slice scan-time differences and 3D head motion using 3 translation and 3 rotation parameters. Subsequently, linear trends and low frequency temporal drifts were removed from the data using a high-pass filter, removing temporal frequencies below 4 cycles per run. After the pre-processing, functional data were co-registered to the high-resolution anatomical volume and normalized to Talairach space. For each subject, a white-matter surface reconstruction was made for each hemisphere separately.

### Cortex-based alignment: transforming data into the same space

A cortex-based alignment (CBA) procedure (Goebel et al., 2006) was applied as implemented in the BrainVoyager QX2.8 software package. For each left and right hemisphere, each subject’s curvature information was aligned to the group (N=34) average in two main steps. First, during a rigid alignment, each subject’s hemisphere was rotated along three dimensions to best match its curvature pattern to one from a randomly chosen target hemisphere. The lower the variability between the two brains, the better the fit after rotation (Goebel et al., 2006; Frost and Goebel, 2012). Second, a non-rigid CBA was performed. Curvature patterns of each subject were used in four different levels of anatomical detail and each aligned to a dynamically changing group average. Starting from low anatomical detail, each subject’s hemisphere was aligned to a group average out of all subjects. During this process, the group average was dynamically updated to most accurately average all hemispheres (measured using the smallest difference across curvature profiles). This sequence was repeated for all four levels of curvature detail until the group average was updated based on the highest level of anatomical detail per subject. During the alignment, we derived (1) a group average hemisphere for the left and right hemispheres, respectively, as well as (2) a transformation matrix that maps between a single vertex of a single-subject cortical surface and a single vertex of a group average cortical surface.

### General Linear Model (GLM)

For the present study, we estimated the mean response to each condition by fitting a general linear model (GLM) to each voxel’s time course for each subject. We then contrasted each condition against the remaining three conditions to obtain a category specific *t*-statistic response for each voxel per run. The resulting *t*-statistic maps were subsequently projected to the cortical surface (integrated across cortical depths along the vertex normals from −1.0 mm to 3.0 mm using linear interpolation). Finally, individual subjects’ cortical surface maps were aligned to the group cortical surface by applying transformation matrices obtained from the CBA procedure.

### Regions of Interest (ROIs)

A ventral temporal cortex (VTC) ROI was defined in each hemisphere on the group average surface described in the CBA section (Fig. 1). The VTC ROI was manually defined by anatomical boundaries (Grill-Spector and Weiner, 2014). Specifically, lateral, posterior, medial, and anterior ROI borders were: the lateral boundary of the occipitotemporal sulcus (OTS), the posterior boundary of the posterior transverse collateral sulcus (ptCoS), the medial boundary of the collateral sulcus (CoS), and the anterior tip of the mid-fusiform sulcus (MFS).

**Figure 1.**
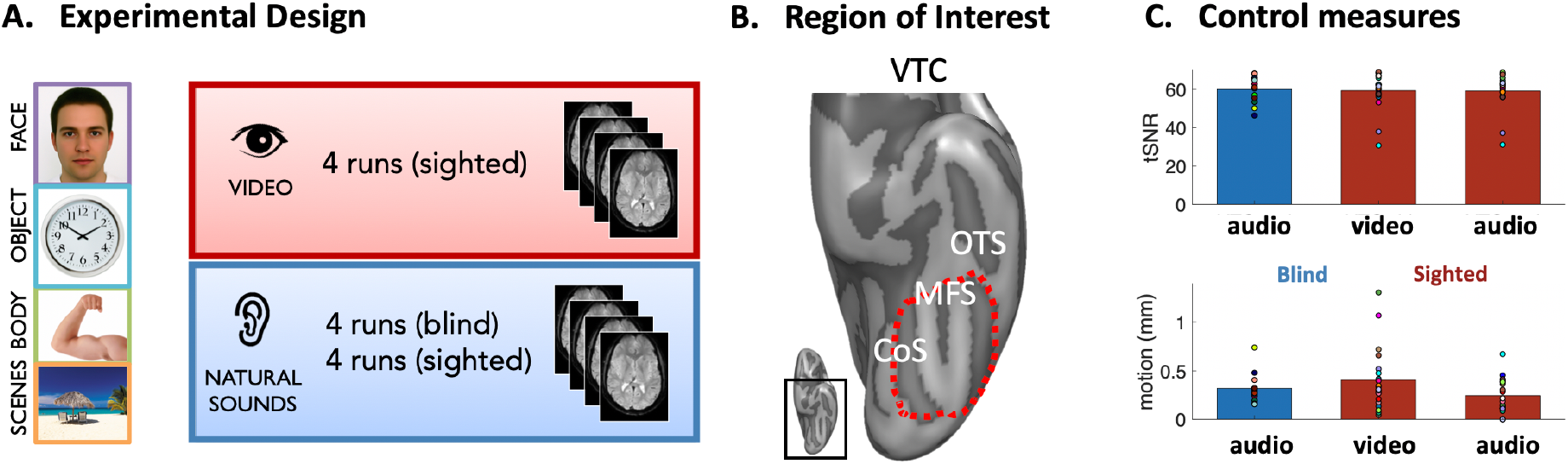
Experimental Design and Regions of Interest. (A) Four different categories (faces, objects, body parts, and scenes) were presented in both visual and auditory modalities. Sighted subjects underwent 4 runs of audio stimuli (first) and 4 runs of video stimuli (second), whereas the blind subjects were presented with 8 auditory runs, of which only the first 4 were used for this study. (B) Using the inflated group average surface (N=34), we manually identified the ventral temporal cortex (VTC) region of interest (ROI) based on macroanatomical landmarks (Materials and Methods). Inset: Ventral view of a left hemisphere in which the black rectangle indicates the zoomed portion depicted in the image at the right. *Dark gray pixels:* sulci; *Light gray pixels:* gyri. *Abbreviations:* A = anterior; P = posterior; M = medial; L = lateral; MFS = mid-fusiform sulcus; CoS = collateral sulcus; OTS = occipito-temporal sulcus. (C) To compare data quality across the two groups, as well as across modalities (video, audio), we computed two quality control measures: (1) the timeseries signal-to-noise ratio (tSNR) across VTC voxels (top row), and (2) motion parameters for each subject and modality for the sighted group (bottom row). Colored dots represent individual subjects. For the sighted (red), the same colored dot across conditions (video, audio) and metrics (tSNR, motion) indicates the same subject. For the blind (blue), the same colored dot across metrics (tSNR, top; motion, bottom) indicates the same subject.

### Representational Similarity Matrices (RSMs)

For each pair of categories, the pattern of selectivity (*t*-statistic map) across all voxels in VTC from the odd runs was correlated with the pattern of selectivity from the even runs. This resulted in a 4×4 representational similarity matrix (RSM) in which a) on-diagonal values correspond to the split-half reliability of the within-category selectivity patterns and b) the off-diagonal entries correspond to the split-half reliability of the between-category selectivity patterns. The RSMs were computed for each individual.

### Permutation testing

We used permutation testing to compare the means of samples without needing to make assumptions about the population distribution. For each analysis, we compared two groups of values: correlations across RSM entries for 1) the sighted video group and 2) the blind group. To establish a null distribution (i.e. the distribution of our data under the hypothesis that there is no difference between the means of the two groups), we randomly shuffled (10,000 times) the group label of each value and computed the difference between the means of the label-permuted groups. This resulted in 10,000 mean differences that are expected under the null hypothesis. Then, we computed the true difference between the correctly labeled groups. To do so, we quantified the proportion, *p,* of permutations in which a label-permuted difference at least as large as the true difference occurred. Because we calculate the mean difference by subtracting the mean of group 2 from the mean of group 1, the resulting *p*-value will be close to a) zero if the mean of group 1 is significantly higher than that of group 2, and b) to 1 when the reverse is true. In cases in which the mean of group 2 is larger, we report 1-*p* to represent the standard of *p* < 0.05 for significant differences and indicate directionality where relevant.

### Statistical analysis: Evaluating RSM group level differences

#### Reliability of category selectivity

In order to evaluate whether a) category selectivity in VTC for each group was equally reliable across split halves of the data (RSM diagonal entries) and b) there were group and category-level differences, we used a 3-way repeated measures analysis of variance (ANOVA) on the magnitude of on-diagonal RSM entries. Specifically, hemisphere (left, right), group (sighted video, blind, sighted audio) and category (face, body, scene, object) were considered as fixed effects, and subjects as a random effect. Post-hoc comparisons were corrected for multiple comparisons using Tukey’s honest significant difference (HSD).

#### Category representation differences between groups

All statistical comparisons only included the off-diagonal entries (called off-diagonal RSM, or odRSM, in further analyses; Ritchie et al., 2017) except for the assessment of test-retest reliability. As the upper and lower triangles within each group are qualitatively similar, we averaged the two halves for further group-level analyses, resulting in 6 off-diagonal entries per subject. To evaluate group level differences for category information (i.e., odRSM), we tested the null hypothesis that there is no difference in odRSM similarity between any pair of subjects across groups. We first generated a null distribution of odRSM-to-odRSM similarity by correlating the odRSMs of randomly drawn subjects to each other, irrespective of the group they belong to. This distribution was computed by selecting subjects with replacement 10,000 times. Next, we estimated the true between-group similarity by taking 10,000 random samples of each subject from the sighted video or audio group (depending on the comparison) and one subject from the blind group. We then calculated the correlation between these two randomly chosen samples, resulting in a distribution of true odRSM similarities. Finally, we used permutation tests as described above to establish whether there was a significant difference between the null distribution of odRSM similarity and the true odRSM similarity distribution. This was repeated for all pairwise comparisons (blind and sighted audio; sighted audio and sighted video), as well as Bonferroni corrected for multiple comparisons.

### Statistical analysis: Quantifying the extent of individual differences

To statistically examine how similar the odRSMs from one member of a group were to odRSMs from other members of that group (e.g. a blind subject’s odRSM to another blind subject’s odRSM), we (1) correlated each subject’s 6-entry odRSM with all other subjects’ odRSMs within the same group, (2) averaged those correlations for that subject, and finally, (3) repeated this for each subject, resulting in one mean correlation to the rest of the group for each subject. This procedure was conducted separately for the blind group, the sighted group (visual stimuli), and the sighted group (audio stimuli; Fig. 4). odRSM similarity was computed as the Pearson’s correlation. While previous studies have implemented alternative metrics that are sensitive only to the rank-ordering of elements (Kriegeskorte et al., 2008), the Pearson’s correlation normalizes for overall activation differences while also remaining sensitive to relative differences across odRSMs. As we are interested in the magnitude of differences across odRSMs, which would be unavailable when using rank order correlations, Pearson’s correlation is more appropriate.

We also evaluated the spread of individual differences within each group. To do so, we used bootstrapping to establish a distribution of sample standard deviations for each group (Fig. 4C) using a four-step procedure. First, for a given group with N subjects (N=20 for sighted visual, 18 for sighted audio, and 14 for blind), we computed the within-group odRSM correlation for each of those N subjects (as described above). Second, we chose N subject’s correlation to the other group members from that group with replacement, such that a given subject’s within-group correlation may be drawn more than once (note that we drew a subject’s correlation to the group rather than the subject’s odRSM). Third, we computed the standard deviation of those N correlation values. This standard deviation provides an estimate of how large the spread of within-group correlations is for our random sample of N subjects. Fourth, we repeated the sampling process 10,000 times, yielding 10,000 sample standard deviations. This analysis was conducted separately for the left and right hemisphere. We then compared the bootstrapped standard deviation distributions of the blind group with the standard deviation distribution of the sighted group, as well as the sighted audio group, using permutation testing as described in *Permutation testing* (Fig. 4). Results were Bonferroni corrected within each hemisphere.

### Statistical analysis: Evaluating RSM similarity between subjects across groups

To investigate how individual subjects contribute to overall similarity or dissimilarity between groups, we asked whether single subjects in a given group were more similar to subjects in their own group or to a subset of subjects in another group, which would support the *individual differences* hypothesis. To test this, we first calculated the correlation between each blind subject’s odRSM and each sighted subject’s 1) audio and 2) video odRSM’s (Fig. 5A). Then, we averaged these two correlations for each blind subject, resulting in a single blind-subject-to-sighted-group similarity estimate for blind-to-sighted-video and blind-to-sighted-audio. For each blind subject, we compared their own-group similarity to the sighted-group similarity by running 10,000 permutations on the own-group and sighted-group correlations in the same manner as described in the section *Permutation testing* (Fig. 5c).

Next, we assessed whether those blind subjects who displayed a high correlation to sighted subjects were also subjects who had high reliability. For all 14 blind subjects, we correlated the subject’s reliability with their similarity to the sighted video group, separately for each hemisphere.

### MRI control measures

A major contributor to between-group differences could always be differences of data quality across groups. To rule out this possibility between groups, we evaluated two data quality measures: timeseries signal-to-noise ratio (tSNR) and head motion.

#### Timeseries signal-to-noise ratio (tSNR) estimation

For each subject and ROI, we a) computed the tSNR by averaging the signal for each voxel within the ROI, b) divided this average by the standard deviation across voxels, and c) calculated an average tSNR across all voxels in the ROI. We then averaged tSNR values between hemispheres. We used a paired-sample t-test to compare tSNR levels between modalities for the sighted group, as well as a two-sample t-test to compare tSNR levels between sighted (video) and blind groups.

#### Motion estimation

For each subject, we computed the root mean square of each of the 6 motion parameters (x, y, z translation; x, y, z rotation) per run, and averaged the results across a) runs and b) motion parameters. We used a paired sample t-test to compare motion estimates within the sighted group across modalities (sighted video/audio), as well as a two-sample t-test to compare motion estimates between sighted (video) and blind groups also across modalities.

## Results

In order to compare the distributed patterns of category selectivity between sighted and blind groups, as well as individual differences within and across each group, we generated 4×4 representational similarity matrices (RSM) for each subject in ventral temporal cortex (VTC). RSMs were calculated separately for each hemisphere and modality (blind: audio; sighted: audio; sighted: video). RSMs are useful for (1) comparing distributed patterns of category information, (2) assessing the discriminability of selectivity for different categories, and (3) assessing reliability of category selectivity (test-retest reliability).

### Similar measurement quality for sighted and blind participants

We first tested if there were differences in measurement quality between blind and sighted subjects during fMRI scanning sessions. We considered two common metrics when comparing groups: <u>head motion</u> and <u>timeseries signal-to-noise</u> ratio (tSNR; Fig. 1C). For head motion, there were no differences between blind and sighted groups (sighted video vs. blind: t(32) = 0.93, p = 0.36; sighted audio vs. blind: t(30) = 0.85, p = 0.40). Similarly, there were also no differences when comparing tSNR between groups (sighted video vs. blind: t(32) = −0.22, p = 0.82; sighted audio vs. blind: t(30) = −0.30, p = 0.76). As video runs were presented after audio runs for each of the sighted subjects, we note that there was a marginal increase in motion for sighted video compared to sighted audio runs that is likely due to run order (sighted audio vs. sighted video runs: t(17) = 1.72, p = 0.10; Fig. 1C). We also note that task performance during fMRI scanning shows that both blind and sighted participants perceived the stimuli in a categorical manner (for details see van den Hurk et al., 2017). These analyses illustrate that both sighted and blind subjects accurately identified categorical boundaries and the data displayed similar MRI measurement quality.

### The reliability of distributed category selectivity in VTC are different between sighted and blind groups

We first compared the split-half reliability of the distributed category-selective patterns between groups. To do so, we visualized the distributed patterns of selectivity to the four categories (faces, scenes, bodies, and objects) across VTC for each subject, separately for independent halves of the data (odd runs/even runs). This allowed us to examine both the spatial layout and reliability of distributed patterns of selectivity for each category compared to all other categories in each sighted subject and each blind subject.

For sighted subjects viewing video stimuli, we found the typical cortical patterns of category selectivity in VTC (Grill-Spector and Weiner, 2014; Weiner et al., 2014; Margalit et al., 2020). This is visible in the representative example sighted subject (Fig. 2A) which shows the typical higher selectivity to faces vs. other stimuli in lateral VTC and the higher selectivity to scenes vs. other stimuli in medial VTC. On the other hand, distributed patterns of selectivity for blind subjects were more variable both within and across subjects. For example, the distributed VTC selectivity to auditory categories in some blind individuals resembled those of the sighted individuals. As shown for the example blind subject in Fig. 2B they show higher selectivity for audio faces in the lateral VTC, and higher selectivity for audio scenes in medial VTC. Like the example sighted subjected in Fig. 2A these patterns of selectivity are similar across odd and even runs. On the other hand, however, some blind individuals exhibited a different cortical topography than the sighted subjects, and for some of these subjects, the reliability of category selectivity varied between odd and even runs as illustrated for the blind subject in Fig. 2C.

**Figure 2.**
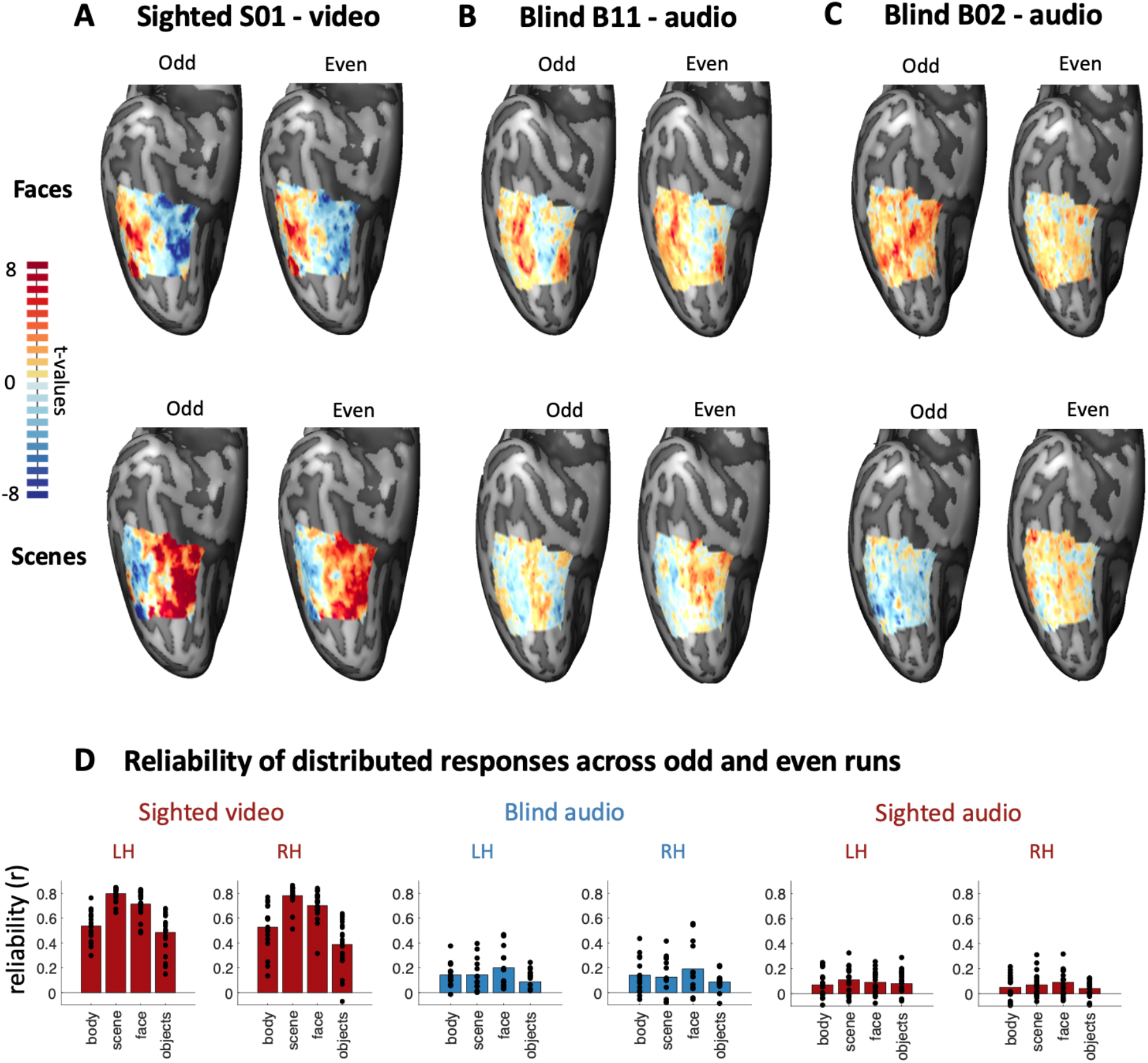
The reliability of distributed category selectivity in VTC is different between sighted and blind groups. (A-C) Unthresholded face-selective (faces vs. all other categories) and scene-selective (scenes vs. all other categories) maps within VTC for odd and even splits of the data. Three example subjects are shown: (A) sighted-visual, (B) blind-audio, and (C) blind-audio. (D) Mean within-category reliability (r) of distributed selectivity for sighted video (left, N=20), blind audio (middle, N=14), and sighted audio (right, N=18). *Dots:* Individual subject reliability. LH: left hemisphere, RH: right hemisphere. Differences in reliability of distributed selectivity in VTC between the sighted and the blind reflect between-group differences irrespective of modality (see Results).

To quantify these observations, we first computed the split-half reliability of distributed category selectivity across VTC and then, compared it between the sighted and blind groups. For sighted visual data (Fig. 2D left column), the average reliability (Pearson correlation between odd/even runs) was between 0.39 and 0.80. For the blind audio data (Fig. 2D middle column), the average reliability was lower, with correlations ranging between 0.09 and 0.20. As these ranges were computed using stimuli from different modalities for each group (video for the former and audio for the latter), as a control, we also compared the reliability of the distributed patterns to auditory stimuli across blind and sighted subjects. We reasoned that if the reliability of the auditory stimuli in the sighted and blind is similar, the differences in reliability between groups may be due to the modality of the stimuli. However, if the reliability between groups still differ for auditory stimuli, it would provide evidence for between-group differences irrespective of the modality of the stimuli.

Our analyses indicate that differences in reliability of distributed patterns of category selectivity in VTC between the sighted and the blind reflect between-group differences irrespective of modality. Specifically, a repeated measures ANOVA with group (sighted visual, blind, sighted audio), category (face, body, scene, object), and hemisphere (left, right) as fixed factors, as well as subject as a random factor, revealed 1) a main effect of group (F(2,391) = 821.30, p = 2e-16), a significant main effect of category (F(3,391) = 37.26, p = 2e-16), 3) a marginal effect of hemisphere (F(1,391) = 3.26, p = .07), and 4) a significant interaction between group and category (F(6,391) = 15.84, p = 2e-16), indicating a difference in the reliability of representations across categories between the two groups (Fig. 2). Post-hoc tests indicate significantly higher reliability for (1) the sighted video group than the blind group (p < 1e-20) and (2) the blind group than the sighted audio group (p = 0.0002). Furthermore, differences in reliability of category selectivity in VTC was not due to differences in tSNR or motion artifacts, as these were statistically indistinguishable across groups (Materials and Methods, Fig. 1C).

### Representational similarity of category selectivity in VTC is different between blind and sighted groups

Qualitatively, average RSMs of category selectivity in VTC in the sighted and blind illustrate on-diagonal values that are higher than off diagonal values (Fig. 3), indicating that distributed patterns of selectivity are more similar within than between categories for both groups. This further indicates that for both sighted video and blind audio at the group level, distributed patterns of selectivity to different categories are distinct (consistent with van den Hurk et al., 2017). We note, however, that the range in values in the sighted video average RSM and the blind audio average RSM are different, in which the former exhibits both higher on-diagonal values and lower off-diagonal values compared to the latter.

**Figure 3.**
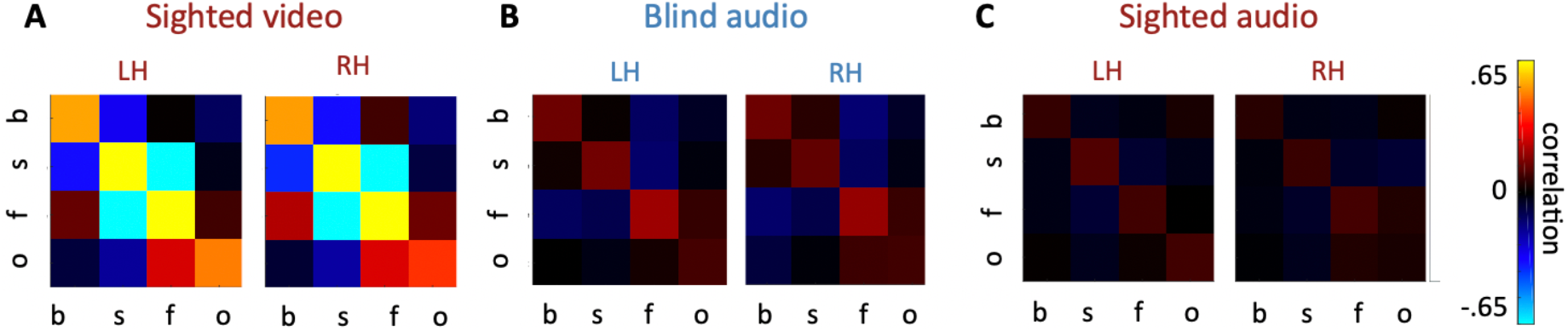
Group averaged Representational Similarity Matrices (RSMs) in VTC differ between sighted and blind groups. *From left to right:* Average RSMs across (A) sighted subjects in response to visual stimuli (N=20), (B) blind subjects in response to auditory stimuli (N=14), and (C) sighted subjects in response to auditory stimuli (N=18). *W*a*rm colors:* positive correlations of distributed category selectivity; *Cool colors:* negative correlations of distributed category selectivity (see colorbar). *Abbreviations:* b = body, s = scene, f = face, o = object, LH = left hemisphere, RH = right hemisphere.

In order to quantitatively assess if there are group differences in VTC RSMs, we first generated a null distribution of similarity (i.e. Pearson correlation) between off-diagonal RSMs (odRSMS) across groups (Materials and Methods). Next, we estimated the true between-group similarity by calculating the pairwise correlations between VTC odRSMs from 10,000 randomly chosen sighted and blind pairs across stimulus modalities. odRSMs from the blind group were significantly less correlated with odRSMs for the sighted video group than expected by chance (p = 1e-04). Additionally, there is a significant difference between blind and sighted auditory participants at the group level (p = 0.04). Together, these results indicate differences in VTC RSMs between sighted and blind groups irrespective of modality.

### Extensive individual differences in representational similarity matrices of VTC category selectivity in the blind

In order to quantitatively compare individual differences in distributed category selectivity in VTC in the blind compared to the sighted, we quantified within-group variability by correlating (Pearson correlation) each subject’s odRSM with all other odRSMs from N-1 subjects, which represents that subject’s similarity to the rest of the group (Fig. 4B). For sighted subjects, the within-group variability was low, indicated by high correlations (r±SD LH: 0.88±0.03; RH: 0.85±0.05), while the within-group variability for blind subjects (r±SD LH: 0.17±0.15; RH: 0.15±0.26) was significantly higher (p < 0.001 in both hemispheres).

**Figure 4.**
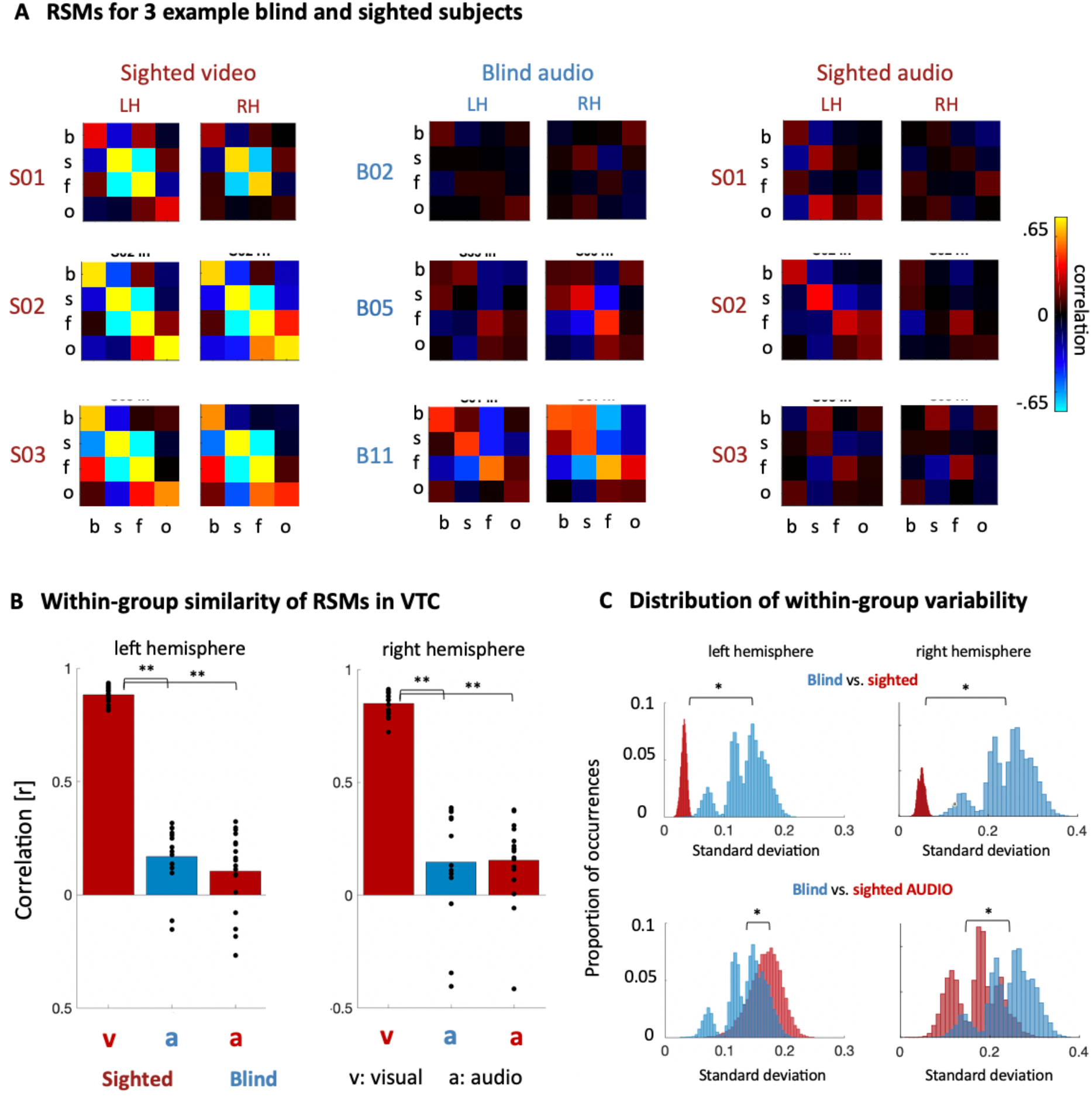
Extensive individual differences in distributed VTC category selectivity in the blind. (A) *From left to right*: RSMs for three example sighted subjects for video stimuli, RSMs for three example blind subjects for audio stimuli, and RSMs for the same example sighted subjects, but for audio stimuli. (B) Average correlation among individual subjects’ odRSM for sighted video (v), blind audio (a, blue), and sighted audio (a, red). Data are plotted separately for the left and right hemisphere. *Dots:* individual subjects; *Asterisks:* p < 0.001. (C) For each group, we bootstrapped (10,000 iterations) subjects and calculated the standard deviation of that iteration, separately for each hemisphere. Histograms in blue refer to the blind group, histograms in red refer to either the sighted visual group (top) or sighted audio group (bottom). *Asterisks*: significantly different (p < 0.05).

Not only were the mean values of the correlations among odRSMs different between the groups, the spread of individual correlations (dots in Fig. 4B) differed between the groups, as the standard deviation of within-group correlations was more widespread in the blind-audio than the sighted-video group. To assess if these differences were significant, we estimated the distribution of the within-group standard deviations using a bootstrapping approach (Materials and Methods, Fig. 4C) and then compared if these distributions differed across the sighted and blind groups. Results showed that the within-group standard deviations were significantly larger in the blind than the sighted group (LH: p = 1e-04; RH: p = 1e-04), as well as further suggest that individual differences in the blind group are not uniformly distributed (see shape of distributions, Fig. 4C). Additionally, our results empirically support that standard deviations were significantly different between the blind-audio group and the sighted-audio group (LH: p = 1e-04; RH: p = 1e-04). Together, these results indicate that (1) there are group differences in VTC representational similarity across the sighted video and blind group, but (2) individual differences in the category representational structure in VTC is more extensive for blind subjects in response to audio stimuli compared to sighted subjects in response to visual stimuli.

### RSMs can be more similar between subjects across groups (sighted and blind) than within group: Further empirical support for the individual differences hypothesis

The prior analyses suggest that individual differences in distributed category selectivity in VTC are more extensive in blind compared to sighted individuals, which supports the *individual differences* hypothesis. Nevertheless, another prediction from this hypothesis is that odRSMs from a subset of blind individuals may be more similar to odRSMs from sighted individuals, while odRSMs from a separate subset of blind individuals may be very different from the sighted. To explore this possibility, we measured the similarity (Pearson correlation) between individual subject odRSMs of each blind subject with each of the sighted subjects (Fig. 5; Materials and Methods).

**Figure 5.**
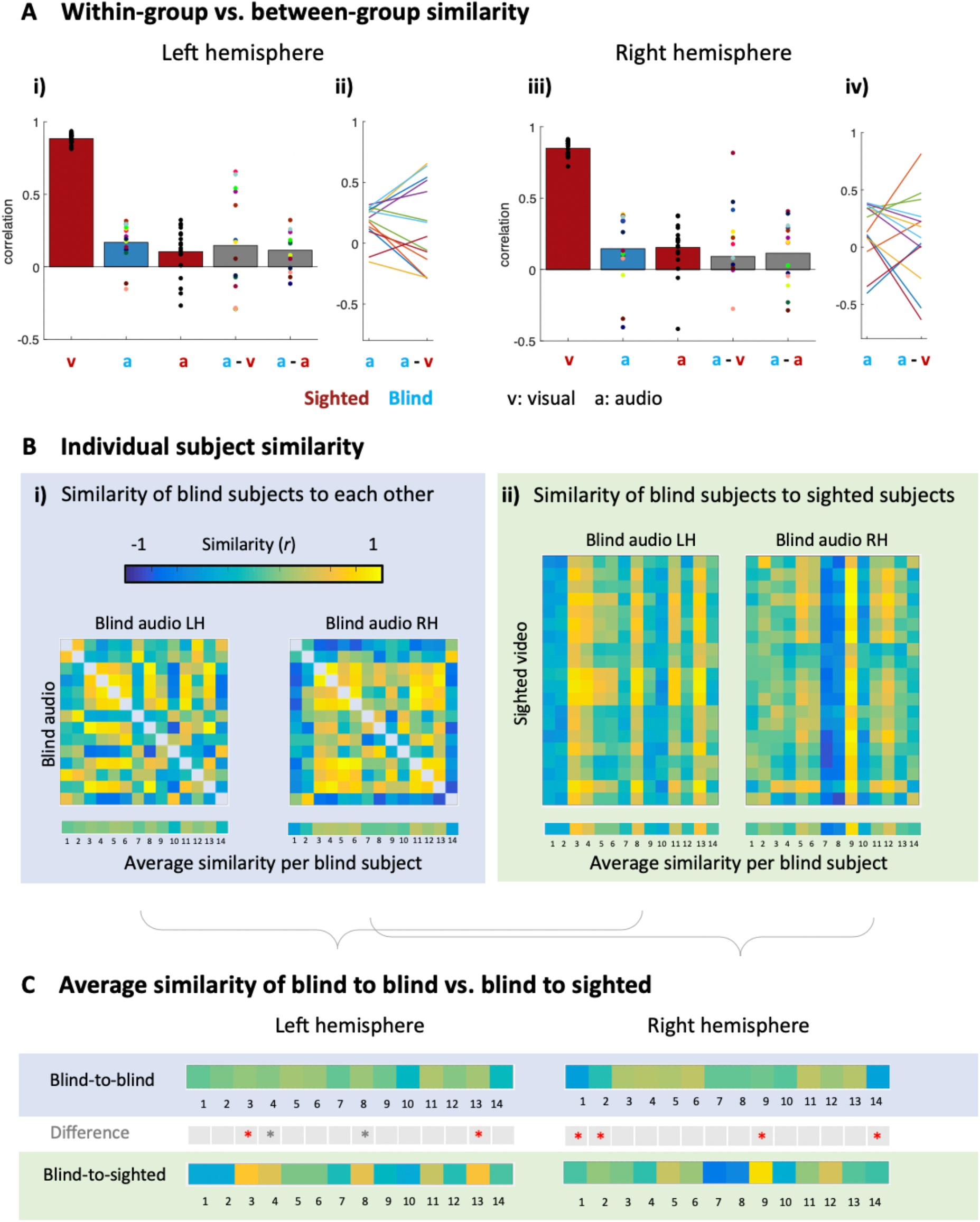
Distributed category selectivity in VTC in a subset of blind subjects is more similar to sighted than blind subjects. (A) *i)* Same organization as Fig. 4A for the first 3 bars. The gray bars represent the between-group similarity: a-v represents the average correlation between a blind subject relative to all sighted subjects in the video condition (individual dots), while a-a represents the average correlation between a blind subject relative to all sighted subjects in the audio condition (individual dots). *ii)* Line graphs depicting correlations between RSMs in individual blind participants (different colors) compared to either other blind participants (a) or other sighted participants (a-v). Lines with positive slopes indicate those participants in which RSMs are more similar to sighted than to blind participants, while lines with negative slopes indicate the opposite. *iii and iv)* Same as i and ii), but for the right hemisphere. (B) (i) Pairwise correlation between the odRSM of each blind subject with each other blind subject, separately for the left and right hemisphere. (ii) The same as (i), but between the odRSM of each blind subject and each sighted control. (C) Average similarity of each blind subject to each blind subject (top row), and to each sighted subject (bottom row). The middle row shows the blind subjects for which odRSMs are more similar to odRSMs from individual sighted subjects than odRSMs from individual blind subjects. Red asterisks: significant difference (ps < 0.05) after False Discovery Rate correction across all 14 comparisons. Gray asterisks: uncorrected significance (ps < 0.05).

Interestingly, this approach revealed that VTC RSMs in a subset of blind individuals are more similar to RSMs in sighted compared to blind individuals (Fig. 5). After performing permutation tests for all 14 comparisons with false discovery rate (FDR) correction, we identified 2 blind subjects in the left hemisphere (B03 p = 0.017, B13, p = 0.047) and 4 blind subjects in the right hemisphere (B01 p = 0.001, B02 p = 0.018, B09 p = 0.001, B14 p = 0.008) that showed a significantly higher correlation to individuals from the sighted group compared to individuals in the blind group (Fig. 5c). In blind subjects who displayed a high correlation to sighted subjects, there was a relationship between reliability and similarity in the left (r = 0.67, p = 0.009), but not the right (r = 0.16, p = 0.40), hemisphere. Intriguingly, this between-group similarity is not due to the ability to perceive light: similarities between individual RSMs can occur between an individual who was born without eyes (B08, B13, B14) and an individual who can see. Together, the combination of our results supports the *individual differences hypothesis* of distributed patterns of category selectivity in the congenitally blind.

## Discussion

Here, we used fMRI to compare distributed category representations (summarized as representational similarity matrices, RSMs) in ventral temporal cortex (VTC) between congenitally blind and sighted subjects and report three main findings. First, there are extensive individual differences in VTC RSMs among blind subjects for audio stimuli, but not sighted subjects for visual stimuli. Second, VTC RSMs in a subset of blind subjects are more similar to RSMs in sighted compared to blind subjects. Third, RSMs in VTC are significantly different between the sighted and congenitally blind at the group level. In the sections below, we discuss the implications of these findings regarding a) the heterogeneity of distributed category representations in VTC of blind subjects, b) how driving factors contributing to the formation of category selectivity in VTC of blind subjects are likely subject-specific, and c) future studies comparing the functional organization of cortical areas between two or more groups.

### Heterogeneity of distributed category representations in VTC of blind subjects

Our results indicate that there are extensive individual differences in distributed category representations within VTC in the blind, and thus, show for the first time that VTC category representations in the blind are more heterogeneous than previously documented. In the context of our own prior work (van den Hurk et al., 2017), the present findings are intriguing as we previously suggested that a) category-selective maps in blind subjects contain core category selectivity that can predict selectivity in the sighted and b) qualitatively, the auditory maps in blind subjects appeared more similar to functional maps based on video stimuli than audio stimuli in sighted subjects. Our results suggest that the latter is not always true when quantitatively comparing VTC category representations from one individual to another between groups. Interestingly, the variability in the blind and also the variability in the sighted for auditory categorical representations in VTC are consistent with what was recently described as “qualitative nuances in the categorical organization of VOTC between modalities [vision and audition] and groups [sighted and blind]” (Mattioni et al., 2020). We build on these qualitative conclusions with our quantifications indicating that there are extensive individual differences of category information in VTC in the congenitally blind.

Our findings of extensive heterogeneity in VTC category representations in blind subjects complement recent studies documenting similarities in the functional organization of visual cortex between the sighted and blind largely at the group level (Mahon et al., 2009; Reich et al., 2011; Striem-Amit et al., 2012a; Kitada et al., 2014). Similar to sighted subjects, blind subjects display large-scale functional maps of animacy (Mahon et al., 2009), as well as functional regions selective for objects (Amedi et al., 2007; Ricciardi et al., 2014), scripts (Reich et al., 2011; Striem-Amit et al., 2012a), tools (Peelen et al., 2014), body parts (Kitada et al., 2014; Striem-Amit and Amedi, 2014), non-manipulable large objects (He et al., 2013), and scenes (Wolbers et al., 2011). Our interpretation of these previous studies is that oftentimes, emphasis is first placed on the detection of a functional activation in a similar anatomical location in VTC of the blind as in the sighted using group analyses, which is then typically followed by a conclusion of similarity between the two groups without quantifying individual differences within and between groups. This approach is consistent with a recent study that included functional maps in VTC of blind individuals at the individual-subject level (Murty et al., 2020).

While these previous findings provide crucial evidence that category-selective representations exist in VTC without visual experience, our findings provide complementary evidence that VTC category representations of blind subjects are extensively variable – sometimes they are correlated with other blind subjects and sometimes they are correlated with sighted subjects. The latter point is critical because while there is a difference between the sighted video and blind groups in the magnitude of the reliability of category representations (Fig. 2), at the individual subject level, this is not *always* true. Together, these findings also have broader implications by indicating that quantitative examinations of individual differences are necessary complements to group analyses in future studies comparing functional representations between blind and sighted groups.

### Driving factors contributing to the formation of category selectivity in VTC of blind subjects are likely subject-specific

To our knowledge, the present findings support a novel hypothesis that driving factors contributing to the formation of VTC category representations in the blind are subject-specific, which complements additional anatomical, functional, and/or experience-dependent factors that may largely generalize across group members (whether blind or sighted) as previously proposed (Tarr and Gauthier, 2000; Malach et al., 2002; Amedi et al., 2007; Martin, 2007; Kalisch et al., 2007; Op de Beeck et al., 2008; Mahon et al., 2009; Kanwisher, 2010; Wolbers et al., 2011; Mahon and Caramazza, 2011; Striem-Amit et al., 2012b; He et al., 2013; Peelen et al., 2014; Ricciardi et al., 2014; Striem-Amit and Amedi, 2014; Grill-Spector and Weiner, 2014; Kitada et al., 2014; Bi et al., 2016; Murty et al., 2020). Specifically, in addition to common mechanisms across blind subjects that contribute to the formation of category representations in the auditory domain within VTC, the way in which VTC, which traditionally processes category representations in a visual domain, forms these representations in blind individuals is likely achieved in a subject-specific manner. This hypothesis is supported by the present data: even the same subtype within the congenitally blind group is not predictive of VTC category representations in a given individual. Specifically, being born without eyes does not predict that category representations in VTC will be similar to or different from VTC category representations in sighted subjects (Fig. 5).

While subject-specific mechanisms likely contribute to the heterogeneity of VTC category representations among congenitally blind individuals, mechanisms that are common across subjects are likely driving the high correspondence and reduced variability of visual category representations in VTC in sighted individuals. We also highlight that there is reduced reliability and extensive individual differences in the sighted for audio category representations in VTC. Thus, it may be the case that functional representations for modalities other than vision in VTC are likely driven by mechanisms that are a) subject-specific and b) independent of being able to see (van den Hurk et al., 2017). Finally, as primary (Aguirre et al., 2016) and multimodal cortical areas (Bedny et al., 2008, 2011) are anatomically and functionally different between sighted and blind individuals, future studies can examine if/how signals from thalamocortical projections or corticocortical connections contribute to individual differences in the functional organization of VTC in the blind.

### Implications for future studies

Our findings have general implications for neuroimaging studies comparing both structural and functional features of cortical areas between two or more groups, especially when within-group homogeneity has not yet been established. For example, while group analyses identify meaningful differences differentiating one group from another, they also mask individual differences within each group. In turn, conclusions from group analyses are often skewed toward assuming homogeneity among members of each group. For example, while in the present study it is generally true that the reliability of VTC category representations is lower in the blind compared to the sighted, reliability of VTC category representations in a subset of blind subjects is comparable to that of the sighted (Fig. 2 Fig. 5). The possibility that group effects could mask effects at the individual subject level is also applicable to any study comparing the functional or structural organization of cortical areas between two or more groups – for example, comparing functional representations between a) children and adults (Cantlon et al., 2006; Golarai et al., 2007; Scherf et al., 2007b; Deen et al., 2016) or b) between neurotypical controls and individuals in a clinical population (Temple et al., 2001; Avidan et al., 2005; Duchaine and Nakayama, 2005; Rykhlevskaia et al., 2009; Green et al., 2019b, 2019a). Future studies can also quantify how different types of stimuli such as soundscapes (Amedi et al., 2007) or haptic substitution (Amedi et al., 2001; Pascual-leone et al., 2005; Reich et al., 2011; Murty et al., 2020) influence individual differences in the blind.

## Conclusion

Using fMRI in congenitally blind and sighted subjects, we empirically show that at the group level, distributed category representations in VTC are more reliable in the sighted (when viewing visual stimuli) compared to the blind (when hearing auditorily-substituted visual stimuli). Despite these group differences, there is extensive heterogeneity in VTC category representations in blind subjects. Together, our findings support a novel idea that driving factors contributing to the formation of VTC category representations in the blind are subject-specific, which complements additional factors that may generalize across group members. More broadly, the present findings caution conclusions of homogeneity across subjects within a group when performing group neuroimaging analyses without explicitly quantifying individual differences.

## Acknowledgements

NSF GRFP (EM)

Excellence of Science grant (EOS; G0E8718N; HUMVISCAT) (HOPdB)

KU Leuven research project C14/16/031 (HOPdB)

NIH R01EY02391501 (KGS)

UC Berkeley start-up funds (KSW)

